# STAAR Workflow: A cloud-based workflow for scalable and reproducible rare variant analysis

**DOI:** 10.1101/2021.09.07.456116

**Authors:** Sheila M. Gaynor, Kenneth E. Westerman, Lea L. Ackovic, Xihao Li, Zilin Li, Alisa K. Manning, Anthony Philippakis, Xihong Lin

## Abstract

**Summary:** We developed the STAAR WDL workflow to facilitate the analysis of rare variants in whole genome sequencing association studies. The open-access STAAR workflow written in the workflow description language (WDL) allows a user to perform rare variant testing for both gene-centric and genetic region approaches, enabling genome-wide, candidate, and conditional analyses. It incorporates functional annotations into the workflow as introduced in the STAAR method in order to boost the rare variant analysis power. This tool was specifically developed and optimized to be implemented on cloud-based platforms such as BioData Catalyst Powered by Terra. It provides easy-to-use functionality for rare variant analysis that can be incorporated into an exhaustive whole genome sequencing analysis pipeline.

**Availability and implementation:** The workflow is freely available from https://dockstore.org/workflows/github.com/sheilagaynor/STAAR_workflow.

## 1 Introduction

The broadening availability of large-scale whole genome sequencing data increasingly permits the analysis of rare variants (RVs), which explain additional heritability and typically have larger effects on complex traits. Modern RV analysis methods, such as the variant-Set Test for Association using Annotation infoRmation (STAAR) method (Li *et al*., 2020), take advantage of variant functional annotations to improve statistical power. However, they can carry heavy computational costs (both memory and runtime) and may not be feasible on local or even some high-performance compute cluster settings for large-scale sequencing studies. Furthermore, frequent updates to computational infrastructure, analysis tools, and genomic data pose obstacles to reproducibility of analysis results. Here, we present STAAR workflow which facilitates the scalable and reproducible analysis of RVs in whole genome sequencing data on cloud platforms.

Our workflow provides a user-friendly, cloud-ready approach to perform set-based testing of RVs using the Workflow Description Language (WDL) (Voss *et al*., 2017). The tool enables genome-wide and candidate RV analyses for both continuous and dichotomous traits while accounting for relatedness and population structure. In contrast to other approaches, it incorporates functional annotations into the RV score-based testing framework. It facilitates gene-centric and genetic region analyses, and was designed to flexibly incorporate aggregation units, annotation types, or variants to condition upon as is frequently performed as follow up. The only required inputs are phenotypic and genotypic data; functional annotations can be obtained from publicly available databases such as FAVOR (Li *et al*., 2020). The tool yields compiled, standard formatted output for use downstream, such as in graphing tools.

Our cloud-based tool is specifically designed to ensure maximally reproducible results and consistent implementation. It uses Docker images to reproduce the computational environment across operating systems without additional software installations or special dependencies, with tracking and versioning on the Dockstore platform (O’Connor *et al*., 2017; Merkel, 2014). The workflow was developed for deployment on NHLBI BioData Catalyst Powered by Terra (NHLBI, 2020; Broad, 2020), but is broadly applicable for any platform that can compile WDL.

## 2 Methods and Implementation

The STAAR workflow operates in three tasks using a generalized linear mixed model (GLMM) approach (Figure 1): null model fitting (Chen *et al*., 2019), score-based testing (Li *et al*., 2020), and results compilation. In all platforms, inputs can be specified in a single, readable JSON object. Analysis methods, computational steps including inputs and outputs with file types, and an example are described in detail in the Supplement.

**Figure 1:**
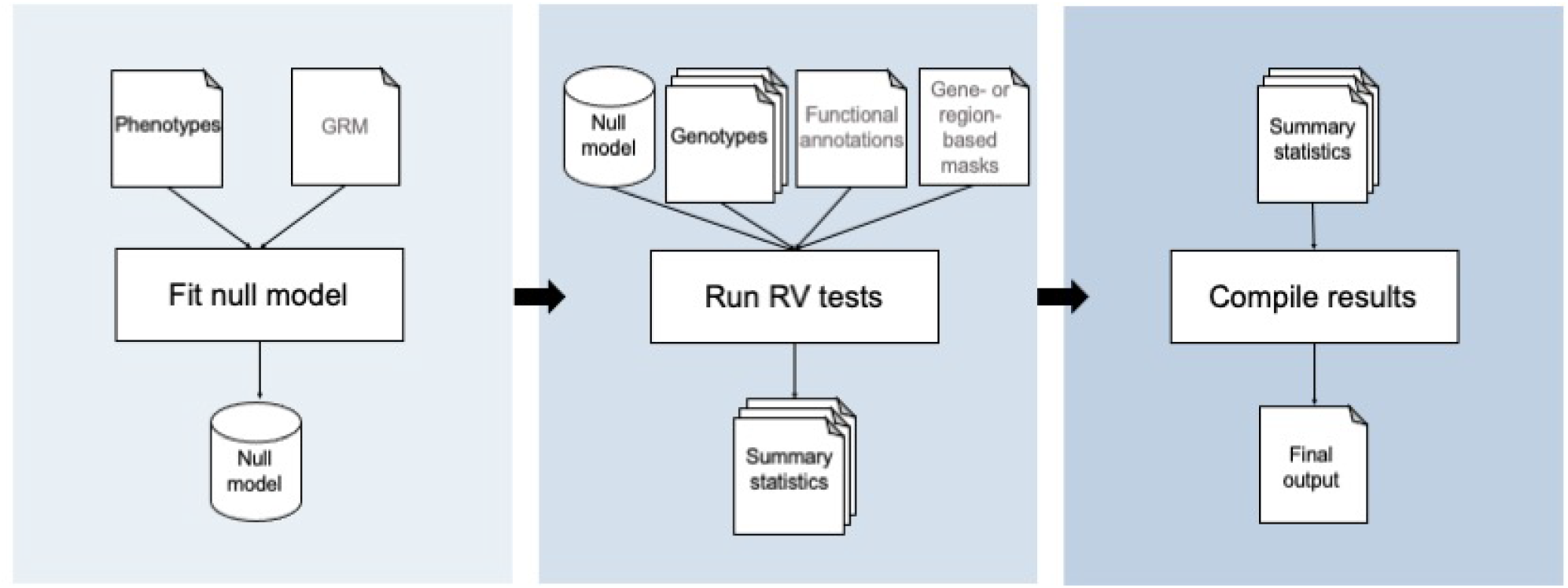
Overview of STAAR WDL workflow. The workflow consists of three sequential tasks. A null model is first fit then passed to the testing task, which performs genetic region or gene-centric RV tests in a parallelized approach (genome-wide or for candidate variant sets). The results of the tests are then aggregated in the final task and output to the user.

The first task generates a null GLMM, a step that can be reused for tests with the same null hypothesis. As input, the null model task only requires phenotypic data. A pre-computed genetic relatedness matrix can be input to account for relatedness via random effects. This task, particularly with large samples and relatedness, is the most memory-intensive. For example, a null model for a quantitative trait with 45,000 samples mimicking the structure of TOPMed can be completed in under three minutes for compute resources with over 100 GB of memory. All tasks use optimized math libraries for efficiency and speed of large matrix operations.

The second task of the workflow performs set-based testing. Given the null model, the only other required file input is genotypic data stored in Genomic Data Structure (GDS) format (Zheng *et al*., 2012; Gogarten *et al*., 2012), as used in large-scale studies like the Trans-Omics for Precision Medicine (TOPMed) program (Taliun *et al*., 2021). For analysis of multiple chromosomes, multiple GDS files can be provided as an array, which are then analyzed in parallel (“scattered”) across separate virtual machines. An essential characteristic of the workflow is the dynamic weighting by functional annotation as introduced in STAAR, thus one can provide functional annotation data as an input file or via variant annotation channels within the GDS. One may further provide aggregation units for gene-centric analysis, a candidate set specification file to limit the analysis to test, and a conditional variant file to specify variants to condition upon with corresponding genotype data. This task uses a custom R script to parse user input and perform score-based tests across the RV sets incorporating a dynamic annotation-based weighting scheme (Li and Li, 2020). Given that RV testing is parallelizable, users can specify the number of cores and partition sets per iteration based on their data and resources. This allows a user to benefit from the on demand resources of the cloud by temporarily utilizing a large amount of resources for rapid completion.

The final task in the workflow compiles the array of result files from the analysis step into a single .txt file for downstream use.

## 3 Conclusion

Used in combination with other GWAS tools, the proposed WDL workflow for RV analysis can facilitate the study of sequencing data to identify novel discoveries and validate findings. In addition to providing a standard suite of RV tests, the main innovation introduced is an efficient cloud-based workflow implementation of the STAAR method that incorporates functional annotations to boost analysis power. The workflow is optimized for out-of-the-box use in cloud ecosystems to support large-scale, reproducible analyses. It is provided in a flexible but automated approach that minimizes investigator effort and accelerates the conduct of analyses. The workflow has been extensively tested on the BioData Catalyst platform and datasets available to the scientific community such as TOPMed. For straight-forward use of STAAR, the presented WDL workflow with documentation is available on Dockstore and GitHub.

## Acknowledgements

The authors thank Beth Sheets and Michael Baumann for their contributions to implementing the workflow on BioData Catalyst.

