## Supplement for "STAAR Workflow: A cloud-based workflow for scalable and reproducible rare variant analysis"

### 1. Statistical model

The STAAR rare variant analysis approach performs aggregate testing of rare variants [1]. The approach extends traditional variant-set tests to incorporate functional annotations into a dynamic weighting scheme. The resulting output is an omnibus test that combines p-values derived across testing frameworks (SKAT, burden, ACAT-V) and functional annotations. The STAAR tests are formulated from a generalized linear model (GLM),

$$g(\mu_i) = \alpha_0 + \boldsymbol{X}_i^T \boldsymbol{\alpha} + \boldsymbol{G}_i^T \boldsymbol{\beta}$$

where $g(\mu)$ is a link function taken to be $g(\mu) = \mu$ for continuous traits and $g(\mu) = \text{logit}(\mu)$ for dichotomous traits. For a set of $p$ variants, for subject $i$ of $n$ individuals consider phenotype $Y_i$ with mean $\mu_i$, $\boldsymbol{X}_i$ the column vector of $q$ covariates, and $\boldsymbol{G}_i$ the column vector of $p$ variants. In the given model, $\alpha_0$ is an intercept, $\boldsymbol{\alpha}$ is a $q \times 1$ vector of coefficients for $\boldsymbol{X}_i$, and $\boldsymbol{\beta}$ is a $p \times 1$ vector of coefficients for $\boldsymbol{G}_i$. This model is extended to a mixed model (GLMM) for data with related samples as

$$g(\mu_i) = \alpha_0 + \boldsymbol{X}_i^T \boldsymbol{\alpha} + \boldsymbol{G}_i^T \boldsymbol{\beta} + b_i,$$

where we assume that $\boldsymbol{b} = (b_1, \dots, b_n)^T \sim N\left(\boldsymbol{0}, \sum_{l=1}^{L} \theta_l \boldsymbol{\Phi}_l\right)$ with variance components $\theta_l$ and known relatedness matrices $\boldsymbol{\Phi}_l$. Additional random effects can also be included. Testing the genotype effects of the $p$ variants, adjusting for covariates and relatedness, has the null hypothesis $H_0: \boldsymbol{\beta} = \boldsymbol{0}$.

In standard score-based testing frameworks, a distributional assumption is made for the genotypic effects $\boldsymbol{\beta}$ to test the null hypothesis. Given the marginal score statistic $S_j = \sum_{i=1}^{n} G_{ij}(Y_i - \hat{\mu}_i)$ for variant $j$ under the null GLM or GLMM, the burden test statistic is given as $Q_{Burden} = \left(\sum_{j=1}^{p} w_j S_j\right)^2$ [2-5], the SKAT test statistic is given as $Q_{SKAT} = \sum_{j=1}^{p} w_j^2 S_j^2$ [6], and the ACAT test statistic is given as $Q_{ACAT-V} = \sum_{j=0}^{p'} w_j^2 \text{MAF}_j(1 - \text{MAF}_j) \tan\left((0.5 - p_j)\pi\right)$ [7]; further details of these conventional tests are provided in the references. In each framework, $w_j$ is a weight defined as a function of the minor allele frequency (MAF) and is commonly taken to be $w_j = Beta(\text{MAF}_j; a_1, a_2)$ where $a_1 = 1$ and $a_2 \in \{1,25\}$.

The STAAR approach extends the weighting scheme by modeling the probability of a variant being causal using functional annotations. Define the effect of variant $j$ on a phenotype as

$$\beta_j = c_j \gamma_j,$$

where $c_j$ is the latent binary indicator of whether variant $j$ is causal, and $\gamma_j$ is the effect size if causal; the effect sizes follow the previously described frameworks, assuming $|\gamma_j| \propto w_j$. Suppose $\pi_j = E(c_j)$ is the probability of variant $j$ being causal, then the effect $\beta_j$ is equivalent to

$$\beta_j = c_j\gamma_j = (1-\pi_j)\delta_0 + \pi_j\gamma_j,$$

where $\delta_0$ is the Dirac delta function indicating that with probability $1-\pi_j$, variant $j$ has no phenotypic association. Then, take $\hat{\pi}_{jk}$ as the probability that variant $j$ is causal considering the $k$th annotation, estimated via the empirical CDF or an indicator of whether the variant is in a qualitative functional group.

Given this incorporation of functional annotations into the effect, the STAAR test statistic is given by extending the standard frameworks as $Q_{Burden,k} = \left(\sum_{j=1}^{p} \hat{\pi}_{jk} w_j S_j\right)^2$ for the burden-based test, $Q_{SKAT,k} = \sum_{j=1}^{p} \hat{\pi}_{jk} w_j^2 S_j^2$ for the SKAT-based test, and $Q_{ACAT-V,k} = \sum_{j=1}^{p} \hat{\pi}_{jk} w_j^2 \mathrm{MAF}_j(1-\mathrm{MAF}_j) \tan\left((0.5-p_j)\pi\right)$ for the ACAT-V-based test for annotation $k$; these have corresponding p-values $p_{Burden,k}, p_{SKAT,k}, p_{ACAT-V,k}$. For functional annotations $k = 0, \dots, K$ the weighted tests are aggregated using the Cauchy combination approach, for instance for the burden-like test we obtain a test statistic $T_{STAAR-B} = \sum_{k=0}^{K} \frac{tan\{(0.5-p_{Burden,k})\pi\}}{K+1}$ with p-value approximated as $p_{STAAR-B} \approx \frac{1}{2} - \frac{\{arctan(T_{STAAR-B})\}}{\pi}$.

To provide a robust test across testing frameworks and weights, the omnibus test statistic STAAR-O is defined as

$$T_{STAAR-O} = \frac{1}{3|\mathcal{A}|} \sum_{(a_1,a_2)\in\mathcal{A}} \left[tan\{(0.5 - p_{STAAR-B(a_1,a_2)})\pi\} + tan\{(0.5 - p_{STAAR-S(a_1,a_2)})\pi\} + tan\{(0.5 - p_{STAAR-A(a_1,a_2)})\pi\}\right],$$

where $\mathcal{A}$ is the set of specified values of $(a_1, a_2)$ as described above to be taken as $\mathcal{A} = \{(1,25), (1,1)\}$. The $P$-value of $T_{STAAR-O}$ is then approximated by

$$p_{STAAR-O} \approx \frac{1}{2} - \frac{\{arctan(T_{STAAR-O})\}}{\pi}.$$

### 2. Computational implementation

The STAAR workflow uses three modularized tasks to perform inference using a generalized linear mixed model approach and provide results. In this approach, a null model assuming no genetic association is first fit and then score-based testing is performed for each RV set. The workflow proceeds in the three tasks: null model fitting, RV testing, and results compilation. Supplementary Table 1 summarizes the inputs and outputs for each step. The workflow was developed on the Terra platform within the BioData Catalyst ecosystem, which allows a simple interactive interface having loaded the workflow from Dockstore [8]. It can also be loaded into other WDL-enabled platforms or used easily on the command line with the Cromwell management system [9].

#### 2.1 Task 1: Null model fitting

In the null model fitting task, phenotypic data is input to fit the previously described model

$$g(\mu_i) = \alpha_0 + \boldsymbol{X}_i^T\boldsymbol{\alpha}$$

for unrelated samples and

$$g(\mu_i) = \alpha_0 + \boldsymbol{X}_i^T\boldsymbol{\alpha} + b_i$$

for related samples, under the null hypothesis $H_0: \boldsymbol{\beta} = \boldsymbol{0}$. In the WDL, this calls the defined task `run_null_model`. The phenotypic data may be limited simply to the outcome provided in a .csv format; the phenotypic data may also include a set of covariates which may include ancestry PCs. The additional required inputs include variable names (outcome, covariate, optional heterogeneous variance group), outcome type (continuous or dichotomous), and output file name as strings. One may optionally provide a pre-computed genetic relatedness matrix to account for sample relatedness in a variety of file formats (.Rds, .Rdata, .csv); a sparse matrix may be provided for scalability. This task uses a custom R script to call the `fit_null_glm` or `fit_null_glmmkin` functions, which are wrapper scripts for the `glm` or `glmmkin` functions of the *stats* package and *GMMAT* packages, respectively. A null model object is generated and returned as an .Rds file.

#### 2.2 Task 2: Rare variant testing

In the testing task, the null model of the first task is required as input; alternatively, a pre-computed null model from previous analyses can be provided. In the workflow, this calls the defined task `run_analysis`. To perform the variant-set tests, genotypic data must be provided in GDS format. The GDS format efficiently compresses and stores genotypes for sequencing studies; it incorporates multiple datasets such as per-sample and per-variant annotations within the object. Large-scale studies currently provide genome-wide sequencing as GDS files. Such files are typically provided split by chromosome, thus this task permits multiple GDS files to be provided in an array to specify analysis across files. They are then analyzed using the scatter function in WDL, which then runs analysis on each task in separate virtual machines in parallel and gathers the output. For the inclusion of functional annotations in the weighting scheme described in STAAR, variant-specific annotation data can be provided as an external file (.Rds, .Rdata, .csv) or by leveraging the variant annotation channels within the GDS and providing annotation locations as input strings. Many repositories and resources exist for compiling relevant functional annotations; they may be general variant annotations such as those defined in CADD [10] or trait- and tissue-specific such as in the Diabetes Epigenome Atlas [11].

In this task, four optional files may be provided to further tailor one's analysis. A file (.Rds, .Rdata, .csv) containing aggregation units can be provided for gene-centric analysis; this provides the opportunity to define sets that are specific to the tested phenotype through trait-specific or tissue-specific sets. Examples of variant set definitions are provided in [1]. A candidate set specification file (.Rds, .Rdata, .csv) can be provided to specify specific aggregation units or windows to limit the analysis to test; by default the workflow analyzes all regions or provided aggregation units. It may be appropriate or necessary to condition on a variant(s) to account for assay effects or determine whether a detected signal is independent of other variants (i.e. GWAS signals) after adjustment. A conditional variant file (.Rds, .Rdata, .csv) can be provided with a corresponding conditional genotype file (GDS) to specify variants to condition upon. Other inputs permit further user specification, including an allele frequency testing threshold, allele count testing threshold, and length of window and overlap for region-based analyses.

The RV testing task uses a custom R script to parse inputs and perform tests calling either the `STAAR` or `STAAR_cond` functions of the STAAR package. Given that RV testing can be fully parallelized, beyond scattering across separate GDS genotype files, users can provide the number of cores and number of iterations to consider splitting sets across. The parallelism is enabled using the *doMC* package, which provides multicore functionality. All output of the testing function is stored, including p-values from standard testing approaches like SKAT and burden (i.e. $p_{Burden,k}$, $p_{SKAT,k}$, $p_{ACAT-V,k}$) for each weight considered, and the resulting omnibus p-value $p_{STAAR-O}$ from the STAAR method.

#### 2.3 Task 3: Result compilation

In the results task, the workflow compiles the array of result files into a single output. This calls the defined task `run_compilation` in the WDL. It takes all results files from the analysis step which are intermediately output as zipped .csv files, and provides a single .txt file for downstream use.

#### 2.4 CWL Implementation

A Common Workflow Language (CWL) version of the tool was also created and can be run on platforms with CWL executors such as NHLBI BioData Catalyst Powered by Seven Bridges.

### 3. Example

We present a simulated example of a rare variant analysis, motivated by the analysis of rare variants by DiCorpo[*] and Gaynor[*] [12], where one significant gene centric signal was identified as associated with fasting glucose. The analysis considered a majority European sample from the TOPMed program (n=26,807). A set of 75 missense RVs in the gene *G6PC2* on chromosome 2 was identified as significant amongst tests of approximately 18,000 aggregate sets of missense rare variants.

#### 3.1 Method

We generated genotypes by simulating 40,000 sequences over 1 Mb regions using the calibration coalescent model (COSI) in order to generate a sample of 20,000 with structure mimicking a European population [13]. We simulated 1,200 distinct regions corresponding to the number of missense sets on chromosome 2, from which we randomly selected regions with lengths of 5 kb within each of the 1 Mb regions to represent genes. The simulated genotypes were prepared then formatted to be stored as a single, chromosome-specific GDS file as data is standardly provided.

Following the result of [12], we simulate phenotypes and annotations assuming a single signal. To leverage the dynamic weighting scheme based on functional annotations as introduced in STAAR, we follow the simulation structure of Li and Li 2021 [1]. We generate $k = 1, \dots, 10$ annotations $A_{j,k}$ for each variant $j$ from a $N(0,1)$ distribution, and store within the GDS file in annotation channels. For the signal 'gene' region, we selected causal variants according to the logistic model

$$\text{logit}\, P(c_j = 1) = \delta_0 + \delta_1 A_{j,1} + \delta_2 A_{j,2} + \delta_3 A_{j,3} + \delta_4 A_{j,4} + \delta_5 A_{j,5},$$

where $\delta_k = \log(5)$. For this signal region, the proportion of causal variants was defined by setting $\delta_0 = \text{logit}(0.18)$ to specify approximately 35% causal variants in the gene. Then, we generated a continuous phenotype for the samples from a GLM according to

$$Y_i = 0.5X_{1i} + 0.5X_{2i} + \beta_1 G_{1j} + \cdots + \beta_s G_{sj} + \epsilon_i$$

where $X_{1i} \sim N(0,1), X_{2i} \sim \text{Bernoulli}(0.5), \epsilon_i \sim N(0,1)$, $G_{mj}$ were the genotypes of the $s$ causal variants in the signal gene with effect sizes $\beta_m$ for $m = 1, \dots, s$. These effect sizes were defined as depending on allele frequency, $\beta_m = c_0 |\log_{10} MAF_m|$ where $c_0 = 0.1$; we assume 80% have positive effects.

#### 3.2 Implementation

Given the data, the pipeline can be run using any of the given specifications. The data as described mimics a user's experience analyzing chromosome 2 for a moderate sample size (n=20,000). The genotype data include 6,927,799 variants, 6,134,932 (88.6%) of which are rare variants (MAF<0.01). Genotypes are stored in the annotated GDS format including 10 simulated annotation PCs (`annot_1-annot_10`) within the GDS file `sim_geno_EU_20K.gds`, which is 1.27 GB in size. There are 1,200 aggregation units stored in `sim_agg_units.csv.gz`, by design, which include 127 rare variants on average. The randomly selected signal region was a set (`simgene_1111`) with 122 rare variants; using the simulation procedure 41 variants were selected to be causal variants and used to generate the phenotype file `sim_pheno.csv`. The average MAF of the signal region was 0.0003 (SD=0.0008).

**Supplementary Table 1**. Workflow inputs and outputs.

| | Required Input | Optional Input | Output |
|---|---|---|---|
| Task 1: Fit null model | • **pheno_file**: [file] file containing the outcome, covariates for the null model (.csv)<br>• **null_file_name**: [string] string containing prefix for | • **covariate_names**: [string] optional, column names in pheno_file of covariate variables, as comma-separated string, to be treated as covariates (string)<br>• **kinship_file**: [file] optional, file containing the kinship matrix for | • **null_file**: [file] file containing output from null model fitting via STAAR (.Rds) |

| | | | |
|---|---|---|---|
| | .Rds output from null model fitting via STAAR (string)<br>• **sample_name**: [string] column name in pheno_file for observation IDs (string)<br>• **outcome_name**: [string] column name in pheno_file for outcome (string)<br>• **outcome_type**: [string] type of variable of outcome, outcome_name in pheno_file, 'continuous' or 'dichotomous' (string)<br>• **null_memory**: [int] requested memory in GB (numeric)<br>• **null_disk**: [int] requested disk size (numeric) | null model with relatedness, row names are sample_names (.Rds, .Rdata, .csv)<br>• **het_var_name**: [string] optional, column name in pheno_file of variable for grouping heteroscedastic errors (string) | |
| Task 2: Run RV tests | • **null_file**: [file] file containing output from null model fitting via STAAR (.Rds)<br>• **geno_file**: [file] file containing genotypes for all individuals from null model, optionally containing the given annotation channels (.gds)<br>• **results_file**: [string] string of name of results file output (string)<br>• **null_memory**: [int] requested memory in GB (numeric)<br>• **null_disk**: [int] requested disk size (numeric) | • **annot_file**: [file] file containing annotations as input with columns 'chr', 'pos', 'ref', 'alt' (.Rds, .Rdata, .csv)<br>• **agds_file**: [string] string indicating whether input geno is an agds file containing the annotations, 'None' [default] (string)<br>• **agds_annot_channels**: [string] comma-separated names of channels in agds to be treated as annotations, 'None' [default] (string)<br>• **agg_file**: [file] file containing the aggregation units in character strings for set-based analysis with columns 'chr', 'pos', 'ref', 'alt', 'group_id' (.Rds, .Rdata, .csv)<br>• **cond_file**: [file] file containing the variants to be conditioned upon with columns 'chr', 'pos', 'ref', 'alt' (.Rds, .Rdata, .csv)<br>• **cond_geno_files**: [file] file containing genotypes for all individuals from null model for conditional analysis; often same as geno_file (.gds)<br>• **cand_file**: [file] file containing units (agg_file required)/windows for candidate sets of interest with columns 'group_id' or 'chr', 'start', 'end' (.Rds, .Rdata, .csv) | • **results**: [file] results file (.csv.gz) |

| | | | |
|---|---|---|---|
| | | • **maf_thres**: [int] AF threshold below which variants will be considered in rare variant analysis, 0.01 [default] (numeric)<br>• **mac_thres**: [int] AC threshold above which variants will be considered in rare variant analysis, 1 [default] (numeric)<br>• **window_length**: [int] length of window for region-based analysis, 2000 [default] (numeric)<br>• **step_length**: [int] length of overlap for region-based analysis, 1000 [default] (numeric)<br>• **num_cores**: [int] number of cores to be used in parallelized analysis, 3 [default] (numeric)<br>• **num_iterations**: [int] number of iterations to run in parallel loop, i.e. how many chunks to split sets into, 20 [default] (numeric) | |
| Task 3: Compile results | *Inputs directly taken from Task 2:*<br>• **results**: [file] results file (.csv.gz) | | • **compiled_results**: [file] compiled results file (.txt) |

Without specification of aggregation units, the workflow defaults to running a genetic region analysis; all inputs, outputs, and corresponding defaults are given in Supplementary Table 1. Given that the data was generated to include the annotations as channels within the GDS file, a chromosome-wide region-based analysis can be outlined for cloud deployment generally in a json file, supposing the data is stored in a Google bucket named `gs://fc-111-222-333`, as:

```
{"STAAR_analysis.agds_annot_channels":"annotation/info/sim_annotation/annot_1,annot
ation/info/sim_annotation/annot_2,annotation/info/sim_annotation/annot_3,annotation
/info/sim_annotation/annot_4,annotation/info/sim_annotation/annot_5,annotation/info
/sim_annotation/annot_6,annotation/info/sim_annotation/annot_7,annotation/info/sim_
annotation/annot_8,annotation/info/sim_annotation/annot_9,annotation/info/sim_annot
ation/annot_10",
"STAAR_analysis.agds_file":"yes",
"STAAR_analysis.covariate_names":"X1,X2",
"STAAR_analysis.geno_files":["gs://fc-111-222-333/sim_geno_EU_20K.gds"],
"STAAR_analysis.null_file_name":"null_model",
"STAAR_analysis.outcome_name":"Y",
"STAAR_analysis.pheno_file":"gs://fc-111-222-333/sim_pheno.csv",
"STAAR_analysis.results_file":"results_regions",
"STAAR_analysis.sample_name":"ID"}
```

In the first task, the null model is generated and output as `null_model.Rds`. The testing and results compilation tasks yielded 247,858 region-based sets, with 74 p-values reported: one for each testing framework (SKAT, burden, ACAT-V) under standard MAF-based weighted ($w_j = Beta\left(\text{MAF}_j; a_1, a_2\right)$ where $a_1 = 1$ and $a_2 \in \{1,25\}$), one for each testing framework weighted by each annotation (1-10), and the STAAR-O omnibus test p-value. The output also provides the chromosome, aggregation unit name or window positions, and number of variants comprising the set. Given the number of tests, there are no sets that meet the Bonferroni-adjusted significance threshold ($0.05/247858 = 2.02 \times 10^{-7}$) as demonstrated in Supplementary Figure 1. The most significant set spans base pairs 154677070-154679069 (48 variants, STAAR-O P-value: $7.27 \times 10^{-6}$).

**Supplementary Figure 1**. Results of genetic region analysis of simulated data. Transformed STAAR-O p-values, resulting from the omnibus test across all testing frameworks and annotations, are plotted at the start of the 2 kb window for the region.

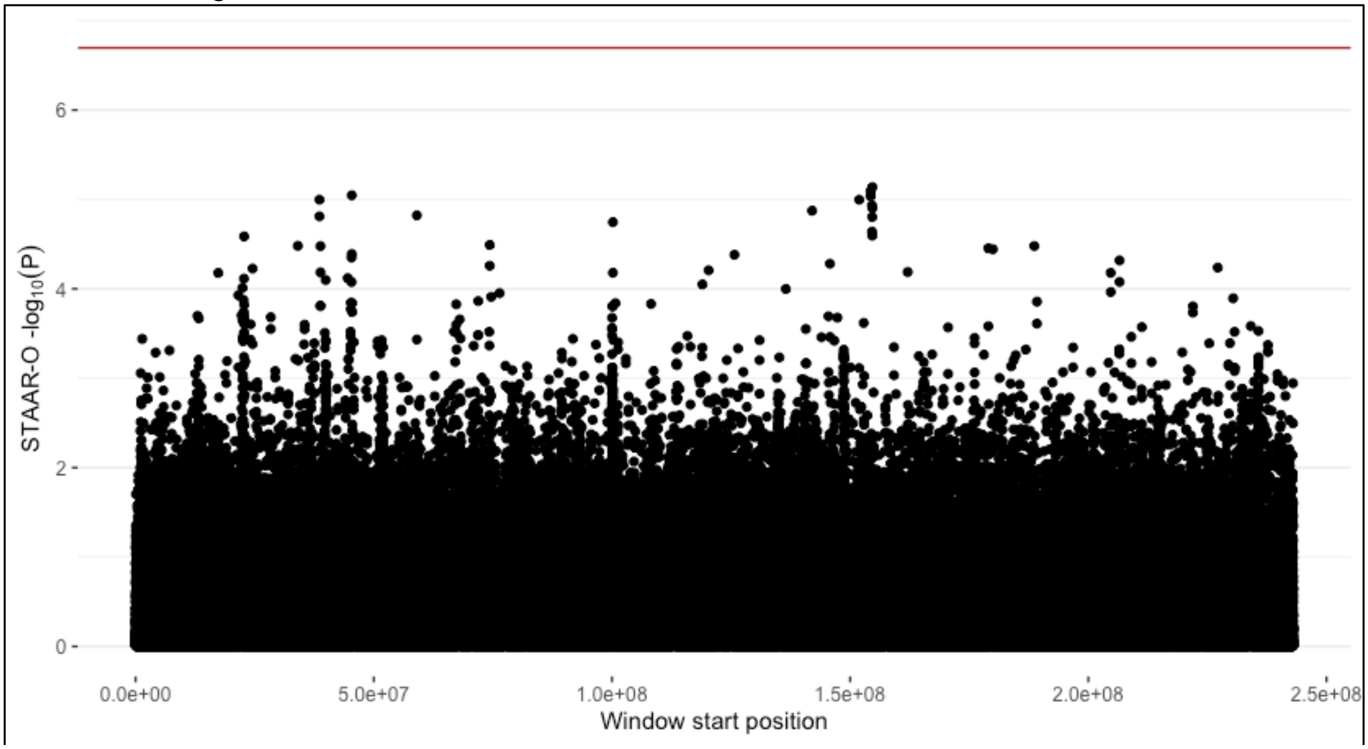


In order to perform the gene centric analysis using the defined aggregation units, the json file can be updated to: (1) include the null model `null_model.Rds` previously computed given that it only needs to be fit once in this score-based testing framework, (2) include the aggregation units `sim_agg_units.csv.gz` defining the 1,200 ‘genes’, and (3) remove the null model arguments:

```
{"STAAR_analysis.agds_annot_channels":"annotation/info/sim_annotation/annot_1,annot
ation/info/sim_annotation/annot_2,annotation/info/sim_annotation/annot_3,annotation
/info/sim_annotation/annot_4,annotation/info/sim_annotation/annot_5,annotation/info
/sim_annotation/annot_6,annotation/info/sim_annotation/annot_7,annotation/info/sim_
annotation/annot_8,annotation/info/sim_annotation/annot_9,annotation/info/sim_annot
ation/annot_10",
"STAAR_analysis.agds_file":"yes",
"STAAR_analysis.agg_file":"gs://fc-111-222-333/sim_agg_units.csv.gz",
"STAAR_analysis.geno_files":["gs://fc-111-222-333/sim_geno_EU_20K.gds"],
"STAAR_analysis.null_file_precompute":" gs://fc-111-222-333/null_model.Rds",
"STAAR_analysis.results_file":"results_genecentric"}
```

In this analysis, only tasks 2 and 3 are run as the null model is reused. A subset of the results of the analysis, the five sets with the smallest STAAR-O p-values, are provided in Supplementary Table 2. The most significant set is `simgene_1111`, as expected by design, with p-value $8.81 \times 10^{-6}$. This test is significant at the Bonferroni-adjusted threshold of $0.05/1200 = 4.17 \times 10^{-5}$ accounting for all gene centric tests performed. Given that all p-values incorporated in the STAAR omnibus p-value are provided in the compiled output, one can investigate the annotations and frameworks driving any significant findings. For this set, the burden tests contributed the most significant p-values.

**Supplementary Table 2**. Results of gene centric analysis of simulated data. STAAR P-values within each of the testing frameworks (SKAT, burden, ACAT-V) considered are provided for the top five sets, ranked by STAAR-O p-value. All STAAR-O p-values incorporate tests using the standard MAF-based weighting scheme with the indicated parameters, as well as tests weighting by each of the 10 simulated annotations.

| Aggregation unit | No. SNV | STAAR-O | STAAR-B (1,25) | STAAR-S (1,1) | STAAR-B (1,25) | STAAR-B (1,1) | STAAR-A (1,25) | STAAR-A (1,1) |
|---|---|---|---|---|---|---|---|---|
| simgene_1111 | 122 | 8.81e-06 | 1.96e-05 | 2.25e-05 | 4.63e-06 | 2.71e-06 | 0.002 | 0.002 |
| simgene_247 | 128 | 2.22e-04 | 6.58e-05 | 8.63e-05 | 1.85e-01 | 1.33e-01 | 0.009 | 0.008 |
| simgene_117 | 123 | 2.84e-03 | 6.89e-02 | 5.05e-02 | 1.08e-03 | 8.77e-04 | 0.116 | 0.092 |
| simgene_93 | 128 | 3.71e-03 | 1.83e-02 | 1.97e-02 | 1.27e-03 | 1.46e-03 | 0.053 | 0.050 |
| simgene_164 | 140 | 4.74e-03 | 2.84e-03 | 2.70e-03 | 2.43e-02 | 1.69e-02 | 0.005 | 0.004 |

To limit these analyses to candidate sets, one can provide a file with either aggregation group names or regions (chromosome, start, end) to analyze. Example files for such candidate analyses are also provided; this can yield cheap, fast, and targeted results. For instance, in the sample gene centric candidate analysis of three simulated genes provided in the workflow distribution (`candidate_gene.csv`) the analysis, including all three tasks, completes at a cost of $0.02 on BioData Catalyst Powered by Terra. The time-cost tradeoff for all analyses, particularly when deployed on cloud platforms, can be optimized based upon one's available resources and timeline. One can further specify their analysis to the desired output by using conditional analysis inputs or adjusting the analysis parameters.

#### 3.3 Availability

The code for generating this sample data is provided in a GitHub repository available at https://github.com/sheilagaynor/STAAR_simulation. A subset of this simulation, defined by the region within a 1Mb window on either side of the signal region, is provided in the workflow distribution as test files (including essential files `genotypes.gds` and `phenotypes.csv`).
